# A randomized, controlled, two-center preclinical trial assessing the efficacy of a new benzodiazepine–dihydropyridine hybrid molecule (JM-20) in rodent models of ischemic stroke

**DOI:** 10.1101/2024.03.08.584085

**Authors:** Jeney Ramírez-Sánchez, André Rex, Sarah McCann, Daniel Schulze, Maylin Wong-Guerra, Luis A Fonseca-Fonseca, Enrique García-Alonso, Ailín Ramírez-Abreu, Ricardo Limonta, Monika Dopatka, Larissa Mosch, Yanier Núñez-Figueredo, Ulrich Dirnagl

## Abstract

JM-20 is a novel multifunctional benzodiazepine molecule with potent neuroprotective effects in rat focal cerebral ischemia. To confirm previous results obtained in single laboratories with small sample sizes, and to provide a robust preclinical evidence base for potential clinical development in stroke, we have performed a two-center preclinical trial with sufficiently large group sizes to detect relevant effects, minimizing biases in experimental design as much as possible (randomization, blinding, predefined in- and exclusion criteria) and increasing external and construct validities by performing experimental focal cerebral ischemia by different surgeons in two different laboratories on two continents, including two species (480 mice and 55 rats), different suppliers, young, young adult, and mature adult animals (range 2 -16 months) as well as comorbid animals (diabetes). While JM-20 improved functional outcomes after middle cerebral artery occlusion in young adult mice at day 7 and appeared to reduce mortality (not statistically significant), it had no effect in mature adult or comorbid (STZ-induced diabetes) mice. Effect sizes, where statistically significant, were modest, and much lower than those reported in the previous studies. Meta-analysis of all individual mouse data did not reveal statistically significant different functional outcomes or mortalities between vehicle- and JM-20-treated animals, although neuroscores and survival were slightly better in JM-20-treated animals. In the less severe model of permanent cortical focal cerebral ischemia in rats, JM-20 significantly reduced brain infarction. We conclude that we were able to confirm the neuroprotective potential of JM-20. However, effect sizes were substantially lower as previously described in small, monocentric trials. Further study is needed to determine whether JM-20 could be effective in less severe cases of focal cerebral ischemia or when used in combination with thrombolysis.

## Introduction

Stroke is frequently associated with severe neurological impairment and mortality. Its incidence is expected to increase due to prolonged life expectancy [1]. Thus far and despite decades of research, the only available pharmacological treatment for the acute phase of cerebral ischemia is tissue plasminogen activator (tPA), applied i.v. within 4.5 h [2].

Due to the complexity of events in cerebral ischemia, it is not realistic to expect that neuroprotective drugs that act on a single pathway or target will have a major impact on outcome. This has prompted a deliberate search for agents that act as multiple ligands or multifunctional drugs [3]. The Center for Pharmaceutical Research and Development (CIDEM, Havana, Cuba) and Havana University (Havana, Cuba) have developed a new multifunctional molecule, JM-20 (3-ethoxycarbonyl-2-methyl-4-(2-nitrophenyl)-4,11-dihydro-1H-pyrido[2,3-b][1,5]benzodiazepine), with promising neuroprotective effect against cerebral ischemia [4–11]. Furthermore, the presence of a positive charge on its benzodiazepine fraction, together with its low molecular weight and high lipophilicity, confers JM-20 physicochemical properties that allow it to cross the blood-brain barrier and reach the mitochondria, facilitated by the negative charge of the organelle matrix.

Previous studies on JM-20 in models related to ischemic brain damage have shown that in mitochondria and synaptosomes isolated from rat brains, JM-20 decreased hydrogen peroxide generation and prevented Ca^2+^ -induced mitochondrial permeability transition pore opening, membrane potential dissipation, and cytochrome c release, all of which are key pathogenic events during stroke [4]. Furthermore, JM-20 inhibited glutamate release from brain synaptosomes and markedly increased glutamate uptake in pure astrocyte cultures as well as co-cultures of neurons and astrocytes [11]. In PC-12 cells JM-20 prevented cell death induced either by glutamate, hydrogen peroxide, or potassium cyanide-mediated chemical hypoxia [5]. In addition, JM-20 protected cerebellar granule neurons from glutamate or glutamate combined with pentylenetetrazole-induced damage at very low micromolar concentrations [5]. In organotypic hippocampal slice cultures exposed to oxygen-glucose deprivation, JM-20 demonstrated neuroprotective effects by modulating neuroinflammation and apoptotic cell signaling pathways [9]. Accordingly, in male Wistar rats subjected to transient middle cerebral artery occlusion (tMCAO, 90 min), oral administration of JM-20 (8 mg/kg) one hour after reperfusion prevented the reduction of activated (anti-apoptotic) Akt levels, neuronal death, and astrocyte reactivity throughout the brain. It also reduced glutamate, aspartate, and GABA increase in cerebrospinal fluid when compared to tMCAO-damaged rats that received vehicle [6, 8]. JM-20 treatment also protected brain mitochondria from ischemic damage, most likely by preventing Ca^2+^ accumulation in the organelle [6]. Remarkably, JM-20 significantly decreased neurological deficit scores (even when administered 8h after reperfusion), edema formation, infarct volume, and histological alterations in different brain regions (cerebral cortex, striatum, and hippocampus) [6]. In a different model of focal cerebral ischemia in rats induced by thermocoagulation of pial blood vessels, JM-20 treatment (4 and 8 mg/kg) significantly decreased asymmetry scores and histological alterations, protecting cortical neurons against permanent focal cerebral ischemia [10].

In summary, we have collected and published substantial evidence that JM-20 has potent neuroprotective effects in rat focal cerebral ischemia, as it dramatically reduced infarct sizes and improved functional outcomes, even when treatment was started many hours after induction of ischemia. However, numerous other promising drugs with similar encouraging preclinical research portfolios failed in subsequent clinical trials. These failures may partially be explained by the fact that preclinical evidence had rested on studies with small sample sizes, as well as questionable internal and low external validity [12]. Under those conditions, false positive results abound, and effects (if true) are grossly overestimated. To provide a robust preclinical evidence base for potential clinical development of JM-20 in stroke, we have performed a two-center preclinical trial with sufficiently large group sizes to detect relevant effects, minimized biases in experimental design as much as possible (randomization, blinding, predefined in- and exclusion criteria) and increased external and construct validities by performing experiments with different surgeons in two different laboratories on two continents, including two species (mouse and rat), different suppliers, young, young adult, and mature adult animals (range 2 -16 months) [13], as well as comorbid animals (diabetes).

## Materials and Methods

### JM-20 treatment

JM-20 was synthesized (quantities sufficient to perform all the procedures) and characterized using techniques and procedures previously described [14]. All treatments were administered orally by gastric gavage (p.o., 10 ml/kg). Immediately before use, JM-20 was suspended in 0.05% carboxymethylcellulose (vehicle). The doses were selected based on previous results [6, 8].

For tMCAO studies, mice received 8 mg/kg JM-20 or vehicle 105 min or 180 min after the occlusion onset and daily for two consecutive days (24 and 48 h after tMCAO). For the permanent focal cerebral ischemia (pFCI) model, rats were treated with 20 mg/kg JM-20 or vehicle 60 min after thermocoagulation and daily until the end of the experiment.

### Focal cerebral ischemia models

All experimental procedures were conducted by trained experimenters following established Standard Operating Procedures (SOPs). tMCAO in mice was induced using an intraluminal filament model, as previously described in detail in a published protocol [15]. Anesthesia was induced with ketamine (75 mg/kg) and xylazine (8 mg/kg) in studies conducted in Havana or with isoflurane in studies conducted in Berlin (1.5-2.0% for induction and 1.0-1.5% for maintenance, with approximately 70/30 N_2_O/O_2_). The left MCA was occluded by introducing a silicon-coated monofilament (7-0 fine MCAO suture Re L910 PK5, Doccol Corp., Sharon MA, USA) into the left internal carotid artery, thereby occluding the MCA origin. After 45 min of MCAO, the animals were re-anesthetized and the filament was removed. During the occlusion period and recovery from surgery, body temperature was maintained using a heating pad or a heated recovery cage. Control animals underwent sham surgery, in which the filament was inserted to occlude the MCA and immediately withdrawn to allow instant reperfusion. After tMCAO, animals had access to soft food and were treated with subcutaneous saline injections to ensure adequate hydration after surgery (once a day until 48 h). Other postoperative measures included pain medication (metamizole, bupivacaine) and a hypercaloric diet for animals losing weight. Further details are given in the preregistered study protocol [22]. Body weight was monitored daily during the first 5 days after tMCAO, and on days 7, 9, and 14 thereafter.

For the induction of permanent focal cerebral ischemia in rats (pFCI), blood thermocoagulation of pial vessels over the motor and sensorimotor cortices was performed, as previously described [10, 16]. Briefly, the animals were anesthetized with sodium pentobarbital (50 mg/kg) and placed in a stereotaxic frame. The skull was surgically exposed, and a craniotomy was performed by exposing the left frontoparietal cortex (+2 to −6 mm anterior-posterior and −2 to −4 mm mediolateral from the Bregma). The blood inside the pial vessels was thermocoagulated transdurally by the apposition of a hot probe close to the dura mater. After the procedure, the skin was sutured and the body temperature was maintained using a heating pad until recovery from anesthesia. Sham-operated control rats were submitted to the craniotomy but without thermocoagulation of pial vessels.

### Animals and husbandry

Animal housing, care, and experimental procedures were executed in accordance with national and institutional guidelines. All procedures were approved by the respective government or institutional committees [Berlin: *Landesamt für Gesundheit und Soziales* (A 0230/18); Havana: Ethics Committee for Animal Experimentation of CIDEM].

Male C57BL/6 mice were supplied by the National Center for the Production of Laboratory Animals (CENPALAB, Cuba; Studies 1, 4, 5) or Janvier Labs (C57BL/6NRj, France, Study 2). Male Wistar rats were supplied by CENPALAB (Cuba, Study 3).

The animals were housed in a temperature-controlled environment (22 ± 2 °C) with a 12-hour light/dark cycle and access to commercial food and water *ad libitum*. The number of animals per cage, the use of environmental enrichment, and the food type were all determined individually by each research group following standard procedures.

### Postoperative care

The first six hours post-op represent a critical period for the animal, with the first two the most critical, therefore the animals were monitored in a warming chamber during this period and then monitored every two hours (until the 6th hour post-op) after being returned to their well-known home cages. Thereafter, the animals were monitored at least twice daily. The cages of the animals were additionally provided with nesting material and a tunnel that remained there until the end of the experiment. The animals received additional moist and mashed food and water in petri dishes on the floor of the cages.

### Exclusion criteria and humane endpoints

Criteria for exclusion from the analysis were: death during surgery or the recovery period (before treatment administration); a neuroscore greater than 6 before surgery (baseline), latency to contact greater than 60 seconds in the baseline ART, or any other signs of neurological alterations before tMCAO induction.

To reduce animal suffering, mice were checked at least two times daily during the first 5 days after surgery. Mice were sacrificed under deep anesthesia if body weight loss exceeded 20% of the initial value, if they permanently circled or were paralytic, or if they had 3 points in the Bederson score [17] or showed other signs of severe neurological impairment. If animals displayed increased defensive reactions, biting, a spasmodically curved back, sunken eyes after fluid loss, or if the abdominal walls were flaccid or painfully taut or the animal could not actively sit up, the animal was also euthanized. Animals with a neuroscore >20 were also euthanized (Study 1, 2, 4, and 5). Additionally, in the comorbidity model (Study 5), mice that died after Streptozotocin (STZ) injection and those with non-fasting blood glucose levels (non-FBGL) less than 250 mg/dl before tMCAO were also excluded.

### Functional outcome tests

Neuroscore (NS) was performed before surgery, 24 h, 72 h, 7, and 14 days after tMCAO or sham-surgery. This test was chosen to evaluate the general status and focal neurologic dysfunction after tMCAO, which induces substantial deficits and was performed as described [18, 19] with some modifications. Summative scores ranged from 0 (no deficits) up to a maximum of 39 points (representing the poorest performance in all items) and were calculated as the sum of the general and focal deficits.

Adhesive removal test (ART) was used to evaluate sensory and motor deficits [20]. A piece of square adhesive tape (2 mm) was placed on the palmar region of the contralateral forepaw. The latency to contact and the latency to remove the adhesive were recorded. Animals were trained for two consecutive days and the baseline test was recorded before tMCAO. In all cases, three trials per animal were performed (with a maximum duration of 120 s), and the average time was calculated. If the mouse failed to remove the adhesive, a value of 121 s was given for that trial.

The corner test (CoT) was performed to assess sensory-motor asymmetry as previously described [21], before tMCAO (baseline) and 7 and 14 days after. Two boards are glued together at an angle of 30° to form a rhombus that creates a gap at the acute angles, which encourages the mice to find the corners. The mouse is placed in the middle of the rhombus and subsequently visits the 30° corners. The mouse moves to the top of the corner, straightens up, and turns in one direction. The number of turns to the left and right are counted and the percentage of right turns of all turns was calculated.

Cylinder test (CyT) was used to evaluate asymmetries in the use of the forelimbs. The test is based on the spontaneous exploratory behavior of rodents, and it reveals forelimb preference when the animal rears to explore the environment. Fourteen days after tMCAO mice were placed in a clear plexiglass cylinder and contacts with the wall during vertical exploration movements with either the left, right, or both forelimbs were counted separately for 5 min. In the pFCI study in rats, a total of 20 contacts were recorded during each trial to prevent habituation to the cylinder. The evaluations were carried out 1 day before and 3, 5, and 7 days after pFCI or sham-surgery. In all cases, the asymmetry score was calculated as previously described [10].

### Infarct volumetry

Infarct volumetry was performed only in study 3 (rat, see below). Rats were transcardially perfused with 0.9% saline and 4% paraformaldehyde in 0.1 mmol/l phosphate buffer. Before proceeding to freeze, the brains were placed in 30% sucrose solution until they reached the bottom of the 50 ml centrifuge tube. Coronal sections of 20 µm were produced using a cryostat, along the entire lesion projected onto the brain surface. The samples were processed for hematoxylin staining and analyzed for quantification of infarct volume. Quantification was performed with ImageJ v1.50 (NIH, USA).

### Streptozotocin-Induced Type 1 Diabetes Model (study 5)

Two weeks before tMCAO surgery, 7-9-weeks-old male C57BL/6N Cenp mice (CENPALAB, Cuba) were fasted for 5-6 hours before a single injection of STZ (120 mg/kg, i.p.). Blood samples were collected by tail-tip clipping and glycemia was measured using a glucometer (SUMASENSOR SXT, Tecnosuma Internacional, Havana, Cuba) according to the manufacturer’s instructions. FBGL were measured before STZ injection and non-FBGL were examined 72 h, 7, and 11 days after to confirm STZ-induced hyperglycemia, and then 1 h after treatment administration (2 h 45 min post-occlusion onset), and 7 and 14 days after tMCAO.

### Study design

We performed 5 substudies, details of which are outlined in Table 1. The experimental design intended to robustly test the reproducibility of results and confirm the efficacy of JM-20 in ischemic stroke models, considering multiple experimental conditions (laboratories from Cuba and Germany), species (mouse and rat), type of focal cerebral ischemia (transient and permanent), age (young and mature adult, ranging 2-16 month) and a comorbid condition commonly found in patients (diabetic hyperglycemia).

**Table 1.**
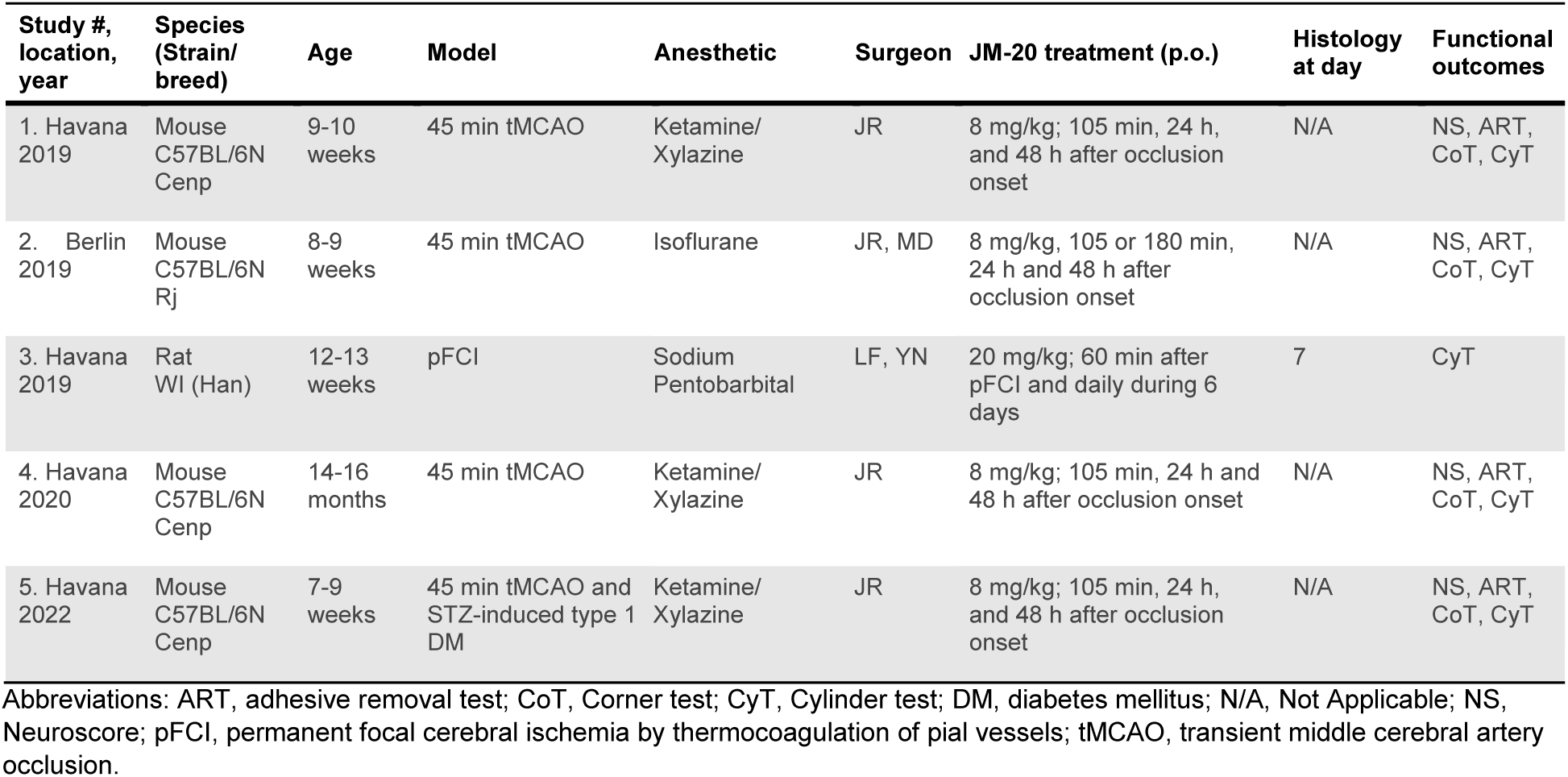
Overview of the five substudies

For tMCAO studies in mice, the predetermined primary endpoint was the NS at day 7. Secondary endpoints were NS at days 1, 3, and 14, as well as sensorimotor and postural asymmetries assessed by ART, CoT, and CyT. Considering the functional consequences of the cortical lesion in the pFCI model in rats (study 3), the primary endpoint was the percentage of asymmetries in the CyT, which has been widely used to assess forelimb asymmetry in this model.

Cuban researchers received training at the Berlin center on surgical procedures for tMCAO and behavioral testing in mice (NS, ART, CoT), and SOPs were drawn up and adapted to local practices and the availability of resources, details of which are described below. Study 2 was preregistered in an Animal Study Registry www.animalstudyregistry.org [22].

The study designs are visualized in Fig. 2a (mouse tMCAO), b (rat pFCI), c (STZ-induced diabetes model), and d (tMCAO in diabetic mice).

**Figure 1.**
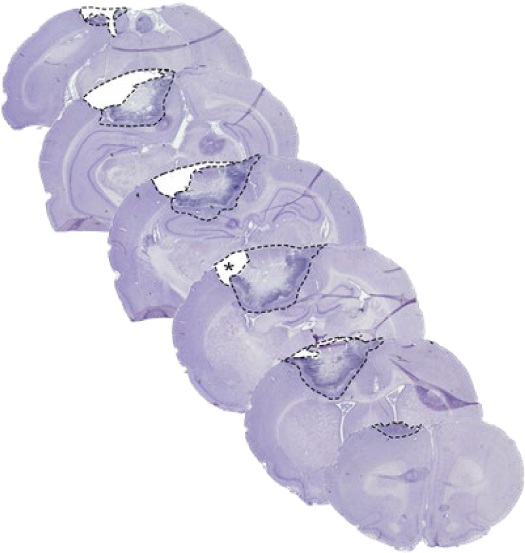
Representative hematoxylin-stained coronal rat brain sections (Study 3). Dashed lines indicate the areas for infarct volumetry, including lost tissue (empty regions, asterisk) surrounding the lesion site 7 days after permanent focal cerebral ischemia (pFCI).

**Figure 2.**
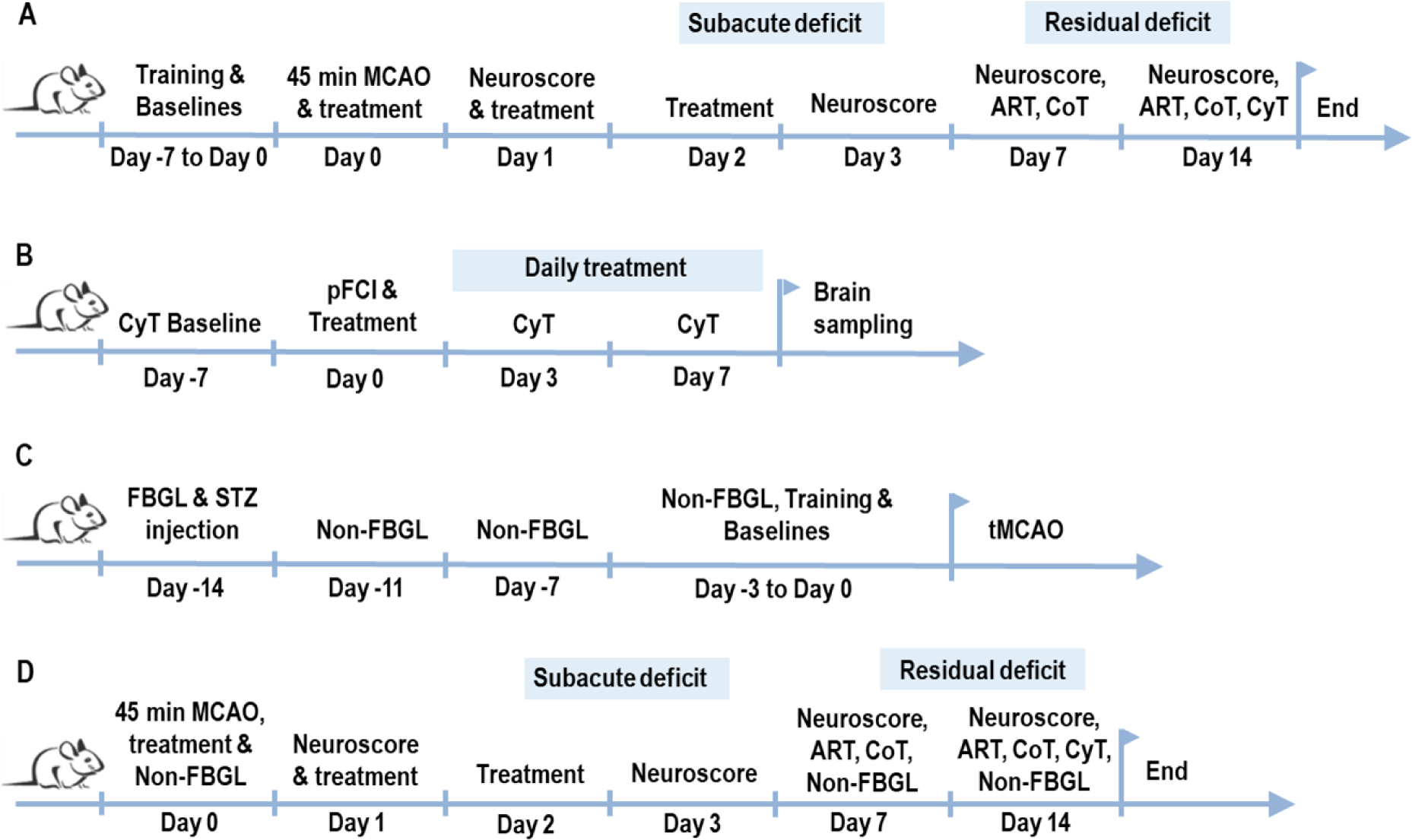
Overview of the study designs. (**A**) mouse tMCAO; (**B**) rat pFCI; (**C**) STZ-induced diabetes model, and (**D**) tMCAO in diabetic mice. ART, adhesive removal test; CoT, Corner test; CyT, Cylinder test; NS, Neuroscore; pFCI, permanent focal cerebral ischemia by thermocoagulation of pial vessels; tMCAO, transient middle cerebral artery occlusion.

### Statistical analysis and methods to prevent bias

The modified intention to treat (ITT) group was the primary analysis population and defined as animals with successful surgery (at least 180 min post-stroke) before the first dose treatment.

Because of high mortality rates in all studies and because we must assume that data were not missing at random (animals were more likely to be missing due to severe stroke), which may lead to attrition biases, we additionally performed secondary analyses using various methods of imputation of missing values (last available NS carried forward, highest possible NS carried forward).

Sample sizes were calculated *a priori* using G*Power software (version 3.1; Heinrich-Heine-Universität Düsseldorf, Düsseldorf, Germany), considering a power of 0.8 and an alpha level of 0.05. For tMCAO models comparing three groups (sham, vehicle, JM-20), assuming an expected effect size of 0.36 (based on previous results in the CoT) and an overall failure rate of 15%, sample sizes of n=31 per group were required; for the study comparing four groups (study 2, additional JM-20 group treated in late time window), an expected effect size of 0.32 was assumed, resulting in n=29 per group. For the pFCI model, comparing three groups (sham, vehicle, JM-20), assuming an expected effect size of 0.58 (based on previous results in the 6-hydroxydopamine model) and an overall failure rate of 15%, sample sizes of n=7 per group were calculated.

The animals were allocated to experimental groups using randomization lists generated online (<u>Randomly assign subjects to treatment groups</u>) on the day of the surgery (after completion of the pre-surgery tests to establish a behavioral baseline). Surgery, behavioral assessments, and analysis of infarct volumes were performed by researchers blinded to the treatment allocation.

All procedures, statistical analyses, and data are reported according to the Animal Research: Reporting of In Vivo Experiments (ARRIVE) guidelines [23]. The complete data set obtained from this study is publicly available at https://zenodo.org/records/10689055.

Data are presented as median with interquartile range or as mean ± standard deviation (Table 2 and Table 3). The normal distribution of all data sets was assessed by the Shapiro-Wilks test. In studies 1, 2, 4, and 5, NS at day 7 (primary outcome) was analyzed using ordinary one-way ANOVA followed by Dunnett’s multiple comparisons test, or unpaired *t*-test or Mann-Whitney U test where indicated. Secondary endpoints were tested using a mixed-effects analysis followed by Dunnett’s multiple comparisons test, corrected for multiple comparisons (body weight gain, blood glucose levels) or ordinary one-way ANOVA followed by Dunnett’s multiple comparisons test (ART, CyT). Mortality was analyzed by comparing the survival curves with the Mantel-Cox log-rank test. In study 3, lesion size and infarct volume were analyzed using the Kruskal-Wallis test followed by Dunn’s multiple comparisons test. Asymmetry was tested using a mixed-effects analysis followed by Dunnett’s multiple comparisons test, corrected for multiple comparisons. All analyses were performed with Prism GraphPad (version 8.0.2 for Windows, GraphPad Software Inc., USA).

**Table 2.** Neuroscore at the defined endpoints and attrition rate (studies 1, 2, 4, 5)

| Study #,<br>location,<br>year | Group | Neuroscore |  |  |  | Total #<br>subjects | Survival<br>(%) |
| --- | --- | --- | --- | --- | --- | --- | --- |
|  |  | 24h | 72h | 1w | 2w |  |  |
| 1. Havana<br>2019 | Sham-Veh | 3.8 ± 1.91 | 4.5 ± 2.58 | 2.3 ± 1.05 | 6.6 ± 2.53 | 20 | 85.0 |
|  | MCAO-Veh | 19.7 ± 7.57 | 11.9 ± 5.52 | 13.8 ± 8.19 | 8.3 ± 2.84 | 38 | 34.0 |
|  | MCAO-JM-20 | 14.5 ± 8.40 | 11.6 ± 6.38 | 6.6 ± 2.53**** | 6.6 ± 2.42 | 36 | 43.0 |
| 2. Berlin<br>2019 | Sham-Veh | 3.0 ± 2.04 | 3.2 ± 1.30 | 2.5 ± 1.44 | 4.0 ± 2.05 | 22 | 95.0 |
|  | MCAO-Veh | 11.3 ± 3.23 | 11.3 ± 3.65 | 9.3 ± 2.55 | 6.6 ± 2.29 | 28 | 54.0 |
|  | MCAO-105 min JM-20 | 11.0 ± 3.41 | 11.3 ± 4.94 | 8.0 ± 1.82 | 7.2 ± 1.30 | 28 | 68.0 |
|  | MCAO-180 min JM-20 | 10.7 ± 4.39 | 10.6 ± 5.40 | 7.0 ± 2.15** | 5.7 ± 1.69 | 29 | 63.0 |
| 4. Havana<br>2020 | Sham-Veh | 7.0 ± 5.34 | 6.5 ± 3.61 | 5.7 ± 3.09 | 5.0 ± 2.30 | 26 | 61.5 |
|  | MCAO-Veh | 21.4 ± 8.02 | 17.7 ± 7.62 | 13.4 ± 2.97 | 11.3 ± 3.22 | 34 | 8.8 |
|  | MCAO-JM-20 | 18.0 ± 7.57 | 13.3 ± 6.76 | 11.1 ± 5.27 | 11.7 ± 5.13 | 34 | 17.6 |
| 5. Havana<br>2022 | Sham-Veh | 5.0 ± 2.70 | 4.6 ± 2.15 | 4.6 ± 1.87 | 4.9 ± 2.37 | 25 | 70.8 |
|  | MCAO-Veh | 13.4 ± 6.28 | 10.0 ± 1.91 | 7.9 ± 3.53 | 8.4 ± 3.43 | 34 | 47.1 |
|  | MCAO-JM-20 | 12.7 ± 7.47 | 9.4 ± 5.46 | 8.1 ± 3.33 | 7.1 ± 2.08 | 32 | 50.0 |

Data are shown as Mean ± Standard Deviation
\*\*\*\* $P < 0.0001$ ; \*\* $P < 0.01$ vs. MCAO-Vehicle group

**Table 3.** Primary and secondary endpoints data and survival rate after permanent focal cerebral ischemia (pFCI) in rats (study 3)

| Study #,<br>location,<br>year | Group | Lesion size<br>(mm<sup>3</sup>) | Infarct<br>volume<br>(mm<sup>3</sup>) | Asymmetry (%) |  |  |  | Total #<br>subjects | Survival<br>(%) |
| --- | --- | --- | --- | --- | --- | --- | --- | --- | --- |
|  |  |  |  | -1d | 3d | 5d | 7d |  |  |
| 3. Havana<br>2019 | Sham-Veh | $1.6 \pm 1.96^{****}$ | $0.1 \pm 0.20^{****}$ | $4.1 \pm 24.23$ | $4.1 \pm 24.23^{**}$ | $2.8 \pm 21.44^{**}$ | $2.8 \pm 21.67^{****}$ | 19 | 100 |
| | pFCI-Veh | $20.2 \pm 12.45$ | $50.1 \pm 28.18$ | $6.8 \pm 10.15$ | $39.9 \pm 32.48$ | $37.3 \pm 40.40$ | $39.3 \pm 24.88$ | 17 | 100 |
| | pFCI-JM-20 | $0.6 \pm 1.23^{****}$ | $0.5 \pm 1.99^{****}$ | $-2.1 \pm 19.02$ | $24.3 \pm 38.63$ | $29.0 \pm 31.56$ | $22.0 \pm 32.86$ | 19 | 100 |

Data are shown as Mean ± Standard Deviation
\*\*\*\* $P < 0.0001$ ; \*\* $P < 0.01$ vs. pFCI-Vehicle group

### Meta-analysis

For evidence synthesis, meta-analysis (not prespecified, but blinded to treatment groups) was conducted using individual data from each mouse study. We compared sham vs MCAO-vehicle vs MCAO-JM-20 as these three groups were present in all four studies (1, 2, 4, and 5; n=342).

For meta-analysis of individual animal data from several studies, statistical modeling has to take the peculiarities of each study into account. We thus used a mixed model approach for analyzing differences between treatments while aggregating across studies. Mixed models adjust for clustering, i.e., animals nested in cages (n=72), which are again nested in studies. Additionally, we allowed the treatment effects to vary between studies. The outcome was NS over the course of 7 days (attrition at day 14 was deemed too high for analysis), as modeled by a log-linear function capturing the initial steep increase, from baseline to post-stroke, followed by a slower decrease (a*time + b*(log(time)), see Fig. 7). Mixed models retain data from cases with missing observations (i.e., attrition) under the assumption that attrition can be explained by other data present, e.g., NS.

Due to the model’s complexity, a Bayesian approach was used for estimation. Thus, 95% credibility intervals (CrI) instead of p-values are reported. Effects were deemed to be significant when the 95%-CrI did not include 0. Attrition over the whole period of 14 days was aggregated using Cox proportional-hazards models stratified for study and cage. All analyses were run in R (version 4.3.1, available at https://zenodo.org/records/10689055) using the packages brms (version 2.20.3) and survival (3.5-5).

## Results

Fig. 3 presents a CONsolidated Standards Of Reporting Trials (CONSORT) style diagram, which demonstrates the flow of all animals enrolled in the study. Total enrollment (animals that were received at study sites and given an identification code) included 480 subjects. Dropouts occurred at various points. Subjects excluded before surgery were considered ineligible. Animals randomized and scheduled for surgery formed the ITT group. Procedural dropouts were subjects excluded due to unsuccessful surgery, while the exclusion from the treatment groups consisted of those that were randomized for stroke surgery but unable to receive the first treatment intervention. The modified ITT group was the primary analysis population and was defined as animals with successful surgery (at least 180 min post-stroke) before the first dose treatment. Partial treatment refers to those animals with successful surgery but which died before treatment completion (during the first 48 h post-surgery). The per protocol group included all subjects who underwent successful surgery, received entire dose treatments, and survived at least 48 h post-surgery. Lost to follow-up were animals who underwent successful surgery, received entire dose treatments but died before the end of the study (14 d post-surgery).

**Figure 3.**
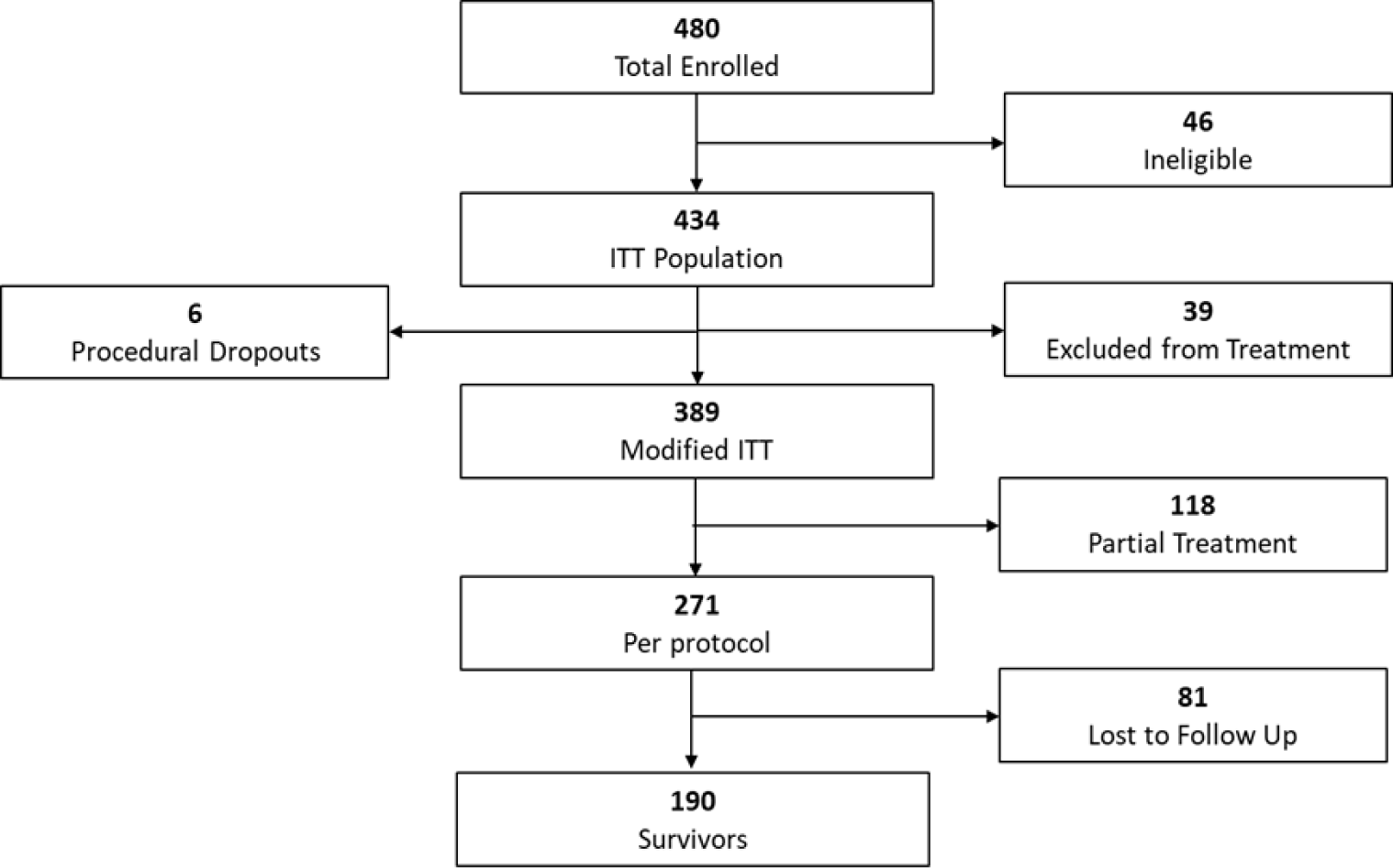
CONSORT style diagram for subjects enrollment and exclusion in tMCAO studies. (Abbreviations in the text).

Table 2 gives a summary of all NS data as well as attrition for substudies 1, 2, 4, and 5. In Study 3 (rat) all animals (n=55) enrolled were eligible and survived until brain sampling on day 7.

Study 1 aimed at evaluating the effect of 8mg/kg JM-20 applied 105 min, 24 h, and 48 h after inducing tMCAO for 45 min in young adult healthy male C57BL/6Cenp mice. The JM-20 treatment group had a 10% higher survival rate than the vehicle treatment group (n.s.).

Compared to vehicle control, JM-20 improved the NS at day 7 (by 52 %, p=0.0001) after tMCAO. The data for study 1 is summarized in Fig. 4A and 5A, and Table 2. No asymmetries were detectable in the CoT, neither in the JM-20 nor the vehicle group. JM-20 and vehicle-treated groups did not differ in the CoT (use of forepaws) or ART (Supplemental Fig. S6). Supplemental Fig. 1A shows only data from animals that survived the entire study (14 days). Supplemental Fig. 2A shows all data points from animals that had died or had to be excluded on or before day 7. Imputations of missing data are shown in Supplemental Fig. 3A and 4A. In Suppl. Fig. 3A missing NS are imputed by assigning the worst possible value (39), while in Suppl. Fig. 4A for missing NS the last available neuroscore is carried forward. Body weight changes (Supplemental Fig. 5A), CoT (Supplemental Fig. 6A), and ART and CyT data are given in Supplemental Table 1.

**Figure 4.**
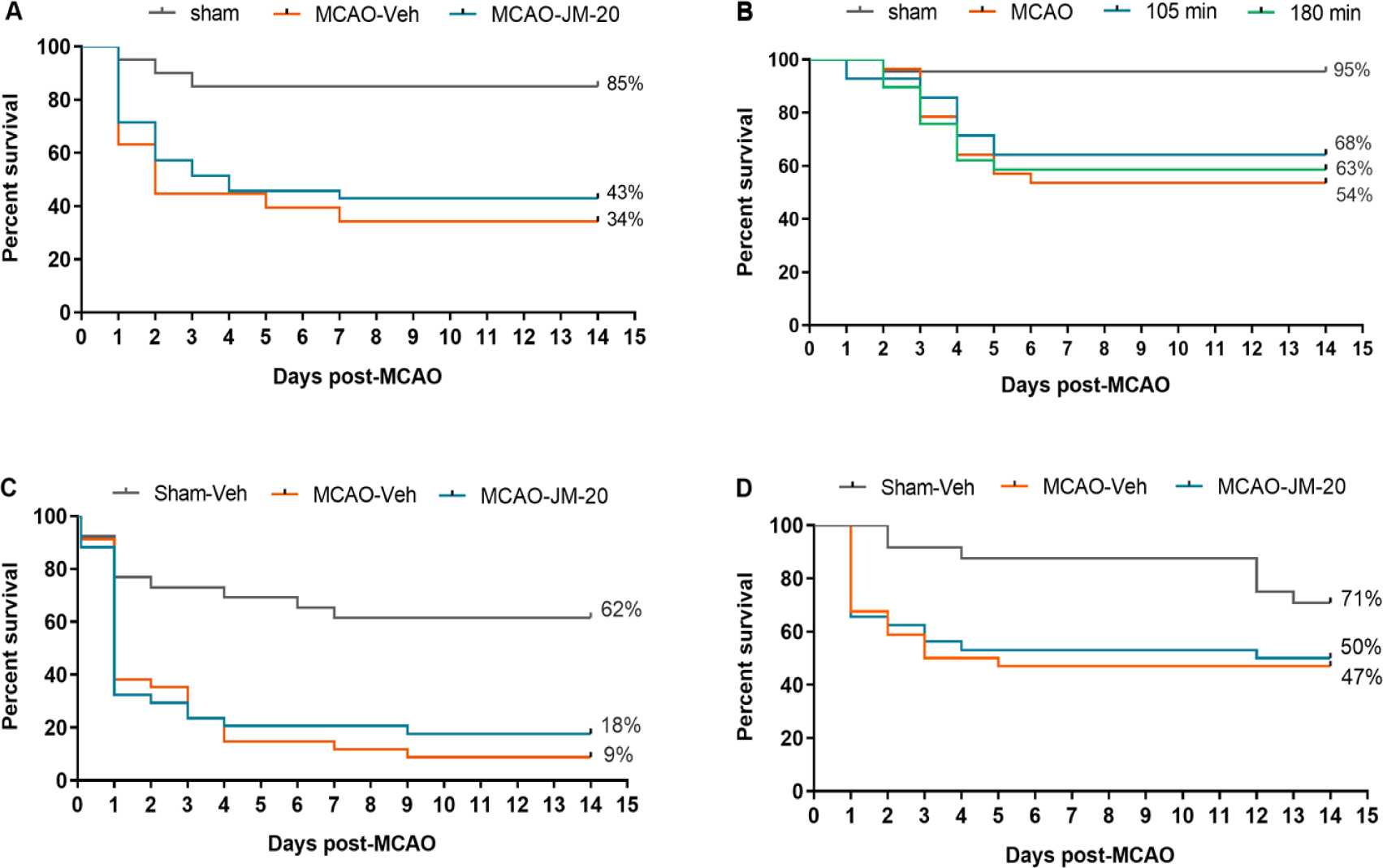
Survival rates. (**A**) Data from Study 1 Havana 2019; (**B**) data from Study 2 Berlin 2019; (**C**) data from Study 4 Havana 2020 and (**D**) data from Study 5 Havana 2022.

**Figure 5.**
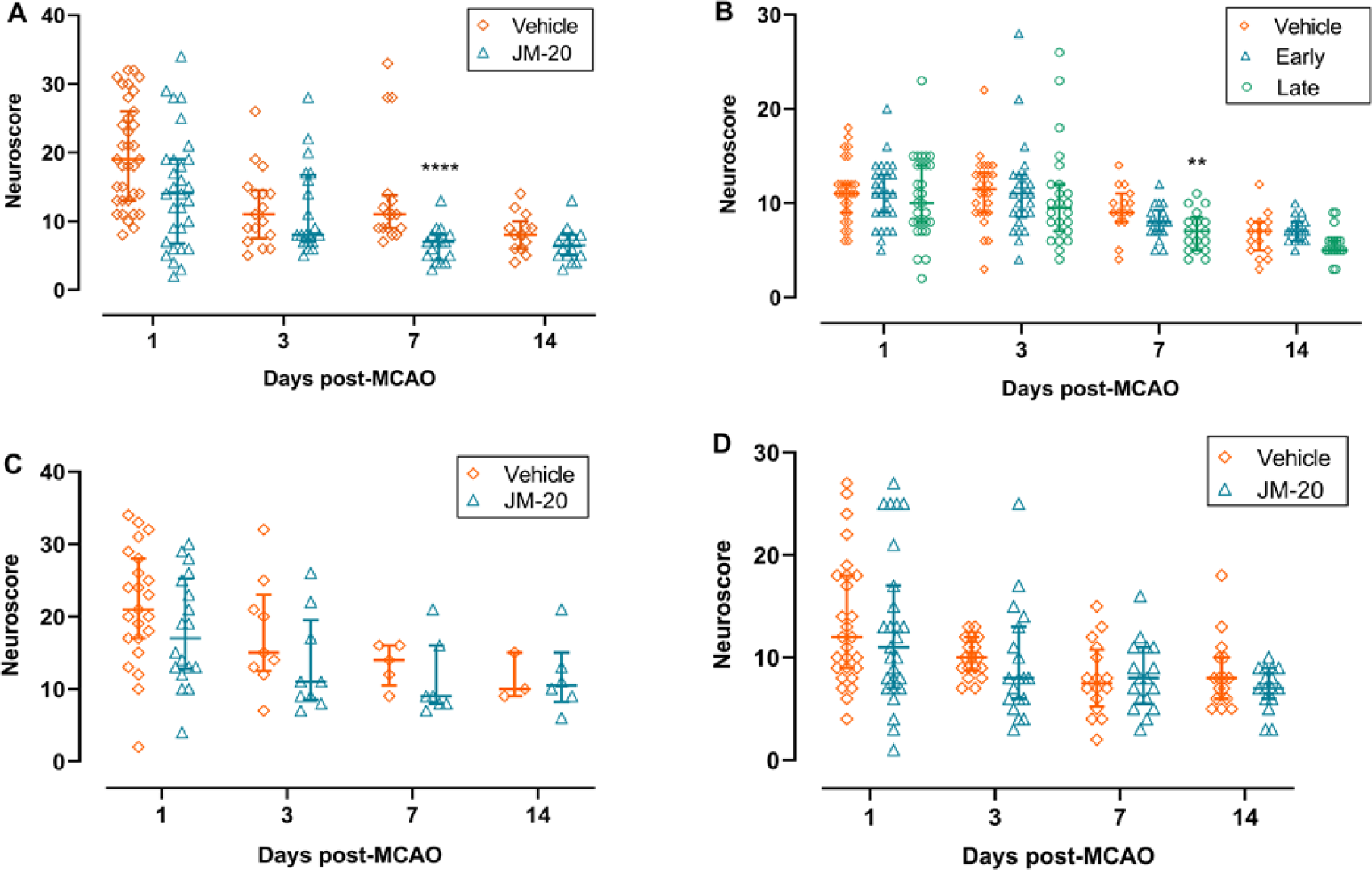
Neuroscore. Median ± Interquartile range. (**A**) Data from Study 1 Havana 2019. \*\*\*\**P*<0.0001 MCAO-Veh vs MCAO-JM-20, by Mann–Whitney U Test. (**B**) Data from Study 2 Berlin 2019. \*\**P*<0.01 MCAO-Veh vs MCAO-JM-20 late treatment, by ordinary one-way ANOVA and Dunnett’s multiple comparisons test. (**C**) Data from Study 4 Havana 2020. *P*>0.05, by Mann–Whitney U Test. (**D**) Data from Study 5 Havana 2022. *P*>0.05, by Unpaired *t*-test.

Study 2 aimed at evaluating the effect of JM-20 (8 mg/kg) in young adult healthy male C57BL/6NRj mice subjected to 45 min tMCAO. JM-20 was given at 105 min, 24 h, and 48 h after the onset of occlusion (‘early time window’) or given at 180 min, 24 h, and 48 h (‘late time window’) after the onset of occlusion.

JM-20-treated groups had a 9-14% higher survival rate than vehicle-treated animals (n.s.). Compared to vehicle control, mice receiving the first JM-20 dose 180 min after onset of occlusion at day 7 had an improved neuroscore (by 25%, p=0.0074). The data for Study 2 is summarized in Fig. 4B and 5B, and Table 2. The late treatment group showed 20 % fewer turns to the left in the CoT, p= 0.046). JM-20 and vehicle-treated groups did not differ in ART (Supplemental Table 1). Supplemental Fig. 1B shows only data from animals that survived the entire study (14 days). Supplemental Fig. 2B shows all data points from animals that had died or had to be excluded on or before day 7. Imputations of missing data are shown in Supplemental Fig. 3B and 4B. In Suppl. Fig. 3B missing NS are imputed by assigning the worst possible value (39), while in Suppl. Fig. 4B for missing NS the last available value is carried forward. Body weight changes are given in Supplemental Fig. 5B, CoT in Supplemental Fig. 6B, and ART and CyT data are given in Supplemental Table 1.

Study 3 aimed at evaluating the effect of JM-20 in pFCI in young adult male Wistar (Han) rats. JM-20 was given at 20 mg/kg, 1 h after thermocoagulation and thereafter daily for 6 days. This substudy attempted to replicate a previously published study [11].

In summary, rats treated with JM-20 showed less asymmetry in the CyT than vehicle-treated animals at all time points tested (n.s.). The data of Study 3 is summarized in Fig. 6 and Table 3. Compared to vehicle-treated rats, JM-20 reduced lost tissue volumes (‘lesion size’) from 20.2 ±12.45 to 0.6 ± 1.23 mm^3^, p< 0.0001 and Infarct volume from 50.1 ± 28.18 mm^3^ to 0.5 ± 1.99 mm^3^, p<0.0001. Body weight changes are given in Supplemental Table 1.

**Figure 6.**
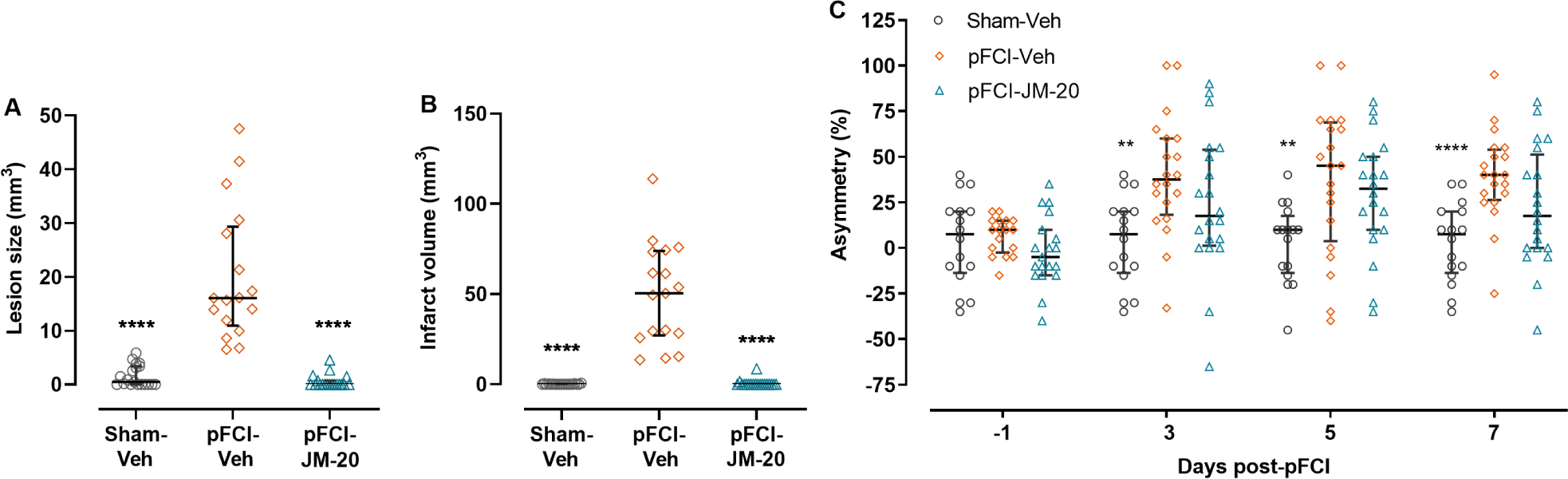
Effect of JM-20 treatment on the ischemic damage induced by permanent focal cerebral ischemia (pFCI) in rats (Study 3). (**A**) The volume of lost tissue 7 days after pFCI. (**B**) Quantitative analysis of the infarcted tissue (pale regions on Hematoxylin-stained coronal sections) surrounding the lesion site. (**C**) Percentage of forelimbs use asymmetry 3, 5, and 7 days after pFCI or sham surgery. No significant differences in forelimb use were detected before damage induction (day -1). Individual values for each experimental subject and median ± Interquartile range are shown. In A and B, ****P<0.0001 *vs* pFCI-Veh group, by Kruskal-Wallis test followed by Dunn’s multiple comparisons test. In C, **P<0.01; ****P<0.0001 *vs* pFCI-Veh group, by Mixed-effects analysis and Dunnett’s multiple comparisons test.

Study 4 aimed at evaluating the effect of JM-20 (8 mg/kg) in mature adult (range 14-16 months) healthy male C57BL/6N Cenp mice subjected to 45 min MCAO when given at 105 min, 24 h, and 48 h after onset of occlusion.

In summary, already in the sham surgery group mature adult mice had a high mortality, which was even higher in mice subjected to tMCAO. The lower NS (day 7) of the JM-20-treated group compared to the vehicle-treated group were not statistically significantly different. Mortality in the JM-20 group was lower than in the vehicle group, but this difference was not statistically significant. The data of study 4 is summarized in Fig. 4C and 5C, and Table 2. Supplemental Fig. 1C shows only data from animals that survived the entire study (14 days). Supplemental Fig. 2C shows all data points from animals that had died or had to be excluded on or before day 7. Imputations of missing data are shown in Supplemental Fig. 3C and 4C. In Suppl. Fig. 3C missing neuroscores are imputed by assigning the worst possible value (39), while in Suppl. Fig. 4C for missing data the last available NS is carried forward. Body weight changes are given in Supplemental Fig. 5C, CoT in Supplemental Fig. 6C, and ART and CyT data are given in Supplemental Table 1.

Study 5 aimed at evaluating the effect of JM-20 (8 mg/kg) in young STZ-induced type 1 diabetic C57BL/6N Cenp mice subjected to 45 min tMCAO. In total, 149 mice were injected with STZ, of which 22 received a second injection. STZ-induced hyperglycemia in 121 mice (>80%) that maintained a non-FBGL with a median close to 400 mg/dl (data shown in Supplemental Fig. 7). Mice were randomly allocated to the experimental groups, with 107 mice receiving tMCAO (excluding mice which did not survive STZ injection, non-hyperglycemic mice or animals showing neurological alterations before tMCAO surgery). Ultimately, 91 mice received treatments (JM-20 or vehicle) and were included in the analysis (excluding those who died during the surgical procedures, due to anesthesia, etc.). Blood glucose levels before and after tMCAO are shown in Supplemental Fig. 7. STZ-induced blood glucose levels after tMCAO were similar in all experimental groups, and JM-20 did not modify them either.

In summary, both vehicle- and JM-20-treated mice had similarly low survival rates (47.1 *vs* 50.0 %). JM-20 treatment had no effect on NS, other functional outcome tests, or mortality. The data of Study 5 is summarized in Fig. 4D and 5D, and Table 2. Supplemental Fig. 1D shows only data from animals that survived the entire study (14 days). Supplemental Fig. 2D shows all data points from animals that had died or had to be excluded on or before day 7. Imputations of missing data are shown in Supplemental Fig. 3D and 4D. In Suppl. Fig. 3D missing NS are imputed by assigning the worst possible value (39), while in Suppl. Fig. 4D, the last available NS is carried forward. Body weight changes are given in Supplemental Fig. 5D, CoT in Supplemental Fig. 6D, and ART and CyT data are given in Supplemental Table 1.

### Results of the meta-analysis of the individual animal data from studies 1, 2, 4, and 5

All treatment effects are reported in comparison with the vehicle-treated tMCAO group. On average, the NS of JM-20 treated animals was 1.40 points lower over time (see Table 4). This averaged meta-analytical benefit was non-significant (95%CrI: -3.99, 1.08). Unsurprisingly, the NS of sham-operated animals was significantly lower than in tMCAO animals (−6.75; 95% Crl:- 9.78, -3.55).

**Table 4.** Mixed model results for aggregating over all four tMCAO studies

| Fixed |  | Random |  |  |
| --- | --- | --- | --- | --- |
| Outcome: NS |  | Study (SD) | Cage (SD) | Mouse (SD) |
| Parallel time courses in treatments |  |  |  |  |
| Intercept | **15.0 (11.0, 18.7)** | **3.53 (1.44, 8.04)** | **1.35 (0.88, 1.85)** | **1.13 (0.19, 1.81)** |
| JM-20 (vs MCAO) | -1.40 (-3.99, 1.08) | **2.69 (0.88, 6.41)** |  |  |
| Sham (vs MCAO) | **-6.75 (-9.78, -3.55)** | **1.87 (0.26, 5.73)** |  |  |
| Day (lin) | **-1.40 (-1.59, -1.22)** |  |  |  |
| Day (log) | **2.49 (2.32, 2.66)** |  |  |  |
| Treatment-dependent time courses |  |  |  |  |
| Intercept | **16.9 (12.9, 20.8)** | **3.60 (1.49, 8.12)** | **1.38 (0.91, 1.87)** | **1.92 (1.44, 2.36)** |
| JM-20 (vs MCAO) | -2.19 (-4.58, 0.23) | **1.86 (0.23, 5.77)** |  |  |
| Sham (vs MCAO) | **-12.1 (-15.8, -8.93)** | **2.75 (0.86, 7.47)** |  |  |
| Day (lin) | **-1.79 (-2.07, -1.52)** |  |  |  |
| Day (log) | **3.33 (3.09, 3.57)** |  |  |  |
| JM-20 x Day (lin) | 0.14 (-0.25, 0.52) |  |  |  |
| Sham x Day (lin) | **1.14 (1.04, 1.80)** |  |  |  |
| JM-20 x Day (log) | **-0.40 (-0.75, -0.05)** |  |  |  |
| Sham x Day (log) | **-2.53 (-2.90, -2.16)** |  |  |  |

Notes: Models take nestedness of measurements over time in each mouse, mice within cages and cages within studies into account. Effects in bold are statistically significant ( $p < .05$ ). Values in parentheses are 95% credibility intervals. SD, standard deviation of fixed effects; tMCAO, transient middle cerebral artery occlusion; NS, Neuroscore.

When allowing different average time courses between the treatment groups, the benefit for JM-20 added up to 1.99 NS points after 7 days (−2.19+.14*7-0.40*(log(7)). This decline was significantly different from the decline in the damaged group treated with vehicle (−0.40, 95%-CrI: -0.75, -0.05) during the steep initial increase in NS in the first 24 h after tMCAO (corresponding to the logistic part of the function, see Fig. 7). However, this finding did not generalize to the long-term development of NS (i.e., the linear part: 0.14, 95%-CrI: -0.25, 0.52). Due to the low NS 24 h after surgery, the decline in NS was significantly different in the sham group (initial increase: -2.53, 95%-CrI: -2.90, -2.16, long-term decrease: 1.14, 95%-CrI: 1.04, 1.80).

**Figure 7:**
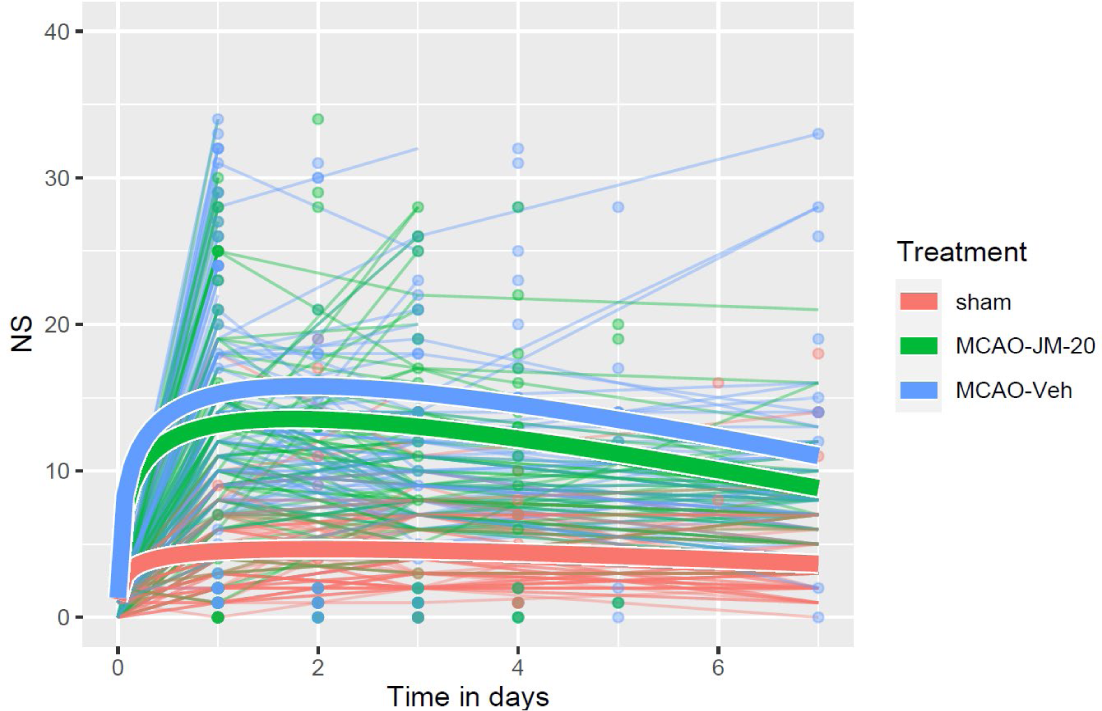
Changes in neuroscore over time per treatment group, showing both individual mice (thin lines) as well as treatment-specific courses as per the log-linear model. (Table 4, second model). Dots indicate an animal’s death.

**Figure 8.**
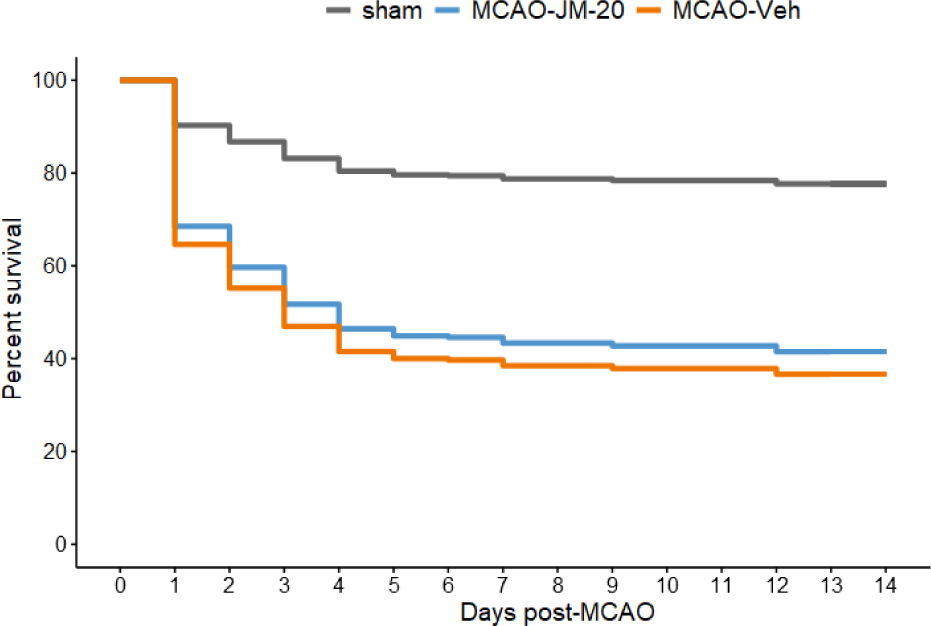
Differences in survival aggregated across the four tMCAO studies (adjusted for study and cage).

In addition, it should be noted that treatment effects varied significantly between studies (second column in Table 4). This finding points towards study-specific heterogeneity in treatment effects. The same held true for the differences between cages and mice. Baseline NS differed significantly between these units of measurement, underscoring their importance in causing variation in effects.

Attrition differed significantly between the sham and the JM-20 groups. For animals treated with JM-20, 14 days non-survival hazard was 5.31 times higher than in the sham group (p <0.001). The same was true for vehicle-treated tMCAO group (HR = 4.75, p <0.001). There was only a small and non-significant difference between JM-20 and vehicle-treated MCAO (HR = 1.12, p = 0.552, after adjustment for study and cage).

## Discussion

A number of exploratory [24] single-center studies have provided evidence that the new multifunctional molecule, JM-20, has potent neuroprotective effects in rat focal cerebral ischemia, even when treatment is started many hours after lesion induction. To investigate whether JM-20 has the potential to be further developed in clinical trials, we have performed a two-center confirmatory preclinical trial [25] with increased statistical power as well as internal and external validity.

Of the 480 mice included (2 mouse strains; transient focal cerebral ischemia model), 389 could be included in the modified ITT population, which was used for primary analysis. On the other hand, 55 rats were included and used for primary analysis in the pFCI model. In studies 1, 2, and 3, the improvement afforded by JM-20 treatment in functional outcome 7 days after lesion, considered the primary endpoint (NS in mice, CyT in rats), reached statistical significance. On secondary analysis, mean neuroscores were more favorable in JM-20-compared to vehicle-treated mice at almost all other measured time points (days 1, 3). A non-prespecified meta-analysis of the individual animal data of all mice subjected to tMCAO revealed a small but statistically significant benefit in the early treatment phase (24 h), which was sustained but did not further increase over the remaining period of 7 days post-tMCAO. In addition, JM-20-treated animals demonstrated significant improvements in some, but not all other functional outcome measurements.

Mortalities, which were substantially higher than anticipated in the sample size calculation (see below), were lower in JM-20-than in vehicle-treated mice. Induction of 45 min MCAO in mature adult mice (study 4) had even higher mortality than in young adult mice, which was lower in JM-20 than in vehicle-treated mice, but this difference was not statistically significant. Likewise, although neuroscores were lower in JM-20-treated animals, statistical improvement in NS at day 7 (the primary endpoint) was not reached in this model. Similarly, induction of diabetes with STZ in young mice (study 5) resulted in a very high mortality, even in sham-operated animals. JM-20 did not improve NS after tMCAO when compared to vehicle control. The mortality rates in diabetic mice treated with JM-20 were not statistically different from those of the vehicle-treated mice. In the pFCI model in rats, no mortality was observed, and JM-20 not only substantially and significantly improved functional outcomes but also reduced infarct sizes compared to the vehicle control, confirming earlier studies in this model [10].

In summary, while JM-20 improved functional outcomes in young adult mice (as assessed by NS) at day 7 and appeared to reduce mortality (not statistically significant), it had no effect in mature adult and comorbid (STZ-induced diabetes) mice. In addition, one has to note that effect sizes, where statistically significant, were modest, and much lower than those reported in the previous studies [6, 8]. Meta-analysis of all individual mouse data did reveal a small but statistically significant improved functional outcome in JM-20-treated animals in the early treatment phase, which was sustained but did not further improve over time. In the less severe model of cortical pFCI in rats (study 3), JM-20 effectively reduced brain infarction.

Our study has a number of weaknesses and limitations. Throughout all experimental groups mortality rates of the tMCAO model were higher than anticipated. The increased mortality was independent of site, surgeon, or strain, and remains unexplained. This has some implications. First, as sample size calculations were based on lower mortality rates, we did not achieve the statistical power we were aiming for. Because post-hoc power analysis is based on the observed effect sizes, it would not add any information beyond the reported p values [26]. We therefore did not set out to determine post-hoc statistical power ‘achieved’ in our study, but do caution that low sample sizes increase type II and type I errors and lead to an overestimation of effect sizes. Second, high mortality rates in vehicle-as well as JM-20-treated mice indicate that we were investigating a rather severe model of ischemic stroke. In elderly patients and those with preexisting health conditions, or when a significant portion of the brain is affected by the ischemic event, clinical stroke can have similarly high mortality rates. However, the efficacy of any treatment is affected by the severity of the disease. In a pathophysiologically complex disease like stroke, it is reasonable to hypothesize that detecting a clinically significant neuroprotective effect becomes increasingly challenging as the severity of tissue damage worsens. The fact that JM-20 was able to improve functional outcomes, albeit modestly, in 3 of the mouse studies, and had a robust effect in a less severe model of pFCI in rats, suggests that JM-20 indeed has neuroprotective potential. However, the strength of the effect may be less pronounced compared to previous, smaller monocentric studies, and restricted to less severe forms of cerebral ischemia. Although not conclusive, our results shed doubt on whether JM-20 neuroprotection extends to individuals of higher age, and with comorbidities.

Most experimental stroke studies are affected by attrition, which in many cases is quite substantial. In the best cases, the number of animals used per group is given, inclusion and exclusion criteria are prespecified, and the number of excluded animals is reported [27]. However, in animal experimentation, data are never missing at random. Animals are more likely to be missing due to severe stroke, resulting in significant selection biases (such as survivor bias) in complete case analysis. This analysis approach, which includes only animals with no missing data for the variable of interest, is considered the de facto standard in preclinical research.

To the best of our knowledge, the only study in experimental research that discusses this important issue and tries to limit its impact is the Stroke Preclinical Assessment Network (SPAN) study [28]. Clinical trials often try to minimize attrition bias by imputation of missing values. The SPAN study, which used a Multi-Arm Multi-Stage (MAMS) statistical design and non-parametric statistics, used multiple imputations assuming the worst rank for missing animals. While for our primary outcome, we used complete case analysis based on the per protocol population, in secondary analyses we have opted for two different, simple imputation methods [29]. In one approach, the last available NS is carried forward, while in the other, a more conservative approach is taken where the worst possible NS (i.e., 39) is imputed for all missing values. Due to the high attrition rates, imputing the worst possible score (39) “clamps” many data points to a single value, resulting in a bimodal distribution with highly unequal variances (see Supplementary Fig. 3). This approach also biases the results towards an endpoint that many of the imputed data points would not have reached. This is because animals that could have potentially survived with lower scores had to be euthanized upon reaching humane endpoints, resulting in the assignment of the highest NS. When the last NS was forwarded (see Supplementary Fig. 4), median neuroscores were lower at all time points in JM-20-treated animals in studies 1 and 2 when compared to the control. While this imputation method in a study like ours may lead to the assignment of more realistic data, we do not know whether this improves the veracity and robustness of our analysis. We conclude regarding the missing data issue that our imputation approaches, while highlighting some of its challenges, did not substantially contribute to improving the robustness of our analysis. Finally, a further limitation in the validity of our study is the use of male animals only. It must therefore remain open whether JM-20 may have differential effects on female mice. We, as well as other researchers, have demonstrated that sex can influence tissue pathophysiology in the context of ischemic stroke [30].

It should be mentioned that a substantial portion of this confirmatory preclinical study, which was executed via the exchange of personnel and material between Cuba and Germany, was overshadowed by the COVID-19 pandemic. Travel restrictions and laboratory shutdowns caused substantial delays, which may partly account for the higher-than-anticipated attrition rate in all experimental groups.

In conclusion, in this international confirmatory, two-center, two-species and two-strain study in focal and permanent cerebral ischemia, designed to increase internal and external validity and sample size compared to the overwhelming majority of experimental stroke studies, we were able to confirm the neuroprotective potential of the multifunctional compound JM-20. However, effect sizes were substantially lower than those previously described in small, monocentric trials. Severe and prolonged ischemia/reperfusion as well as advanced age or comorbidities may even further limit the neuroprotective capability of JM-20. Our results suggest exercising caution when considering the clinical advancement of JM-20 in stroke. Further study is needed to determine whether JM-20 could be effective in less severe cases of focal cerebral ischemia or when used in combination with thrombolysis.

## Acknowledgments

Funded by the German Ministry for Education and Research (BMBF Grant CUB17WTZ-064)

## Data availability statement

All data is available on Zenodo (DOI 10.5281/zenodo.10689054)

## Abbreviations

ART: adhesive removal test
CONSORT: CONsolidated Standards Of Reporting Trials
CoT: Corner test
CyT: cylinder test
FBGL: Fasting blood glucose levels
ITT: intention to treat
NS: neuroscore
pFCI: permanent focal cerebral ischemia
SOPs: Standard Operating Procedures
STZ: Streptozotocin
tMCAO: transient middle cerebral artery occlusion

