## Supplementary materials for "A randomized, controlled, two-center preclinical trial assessing the efficacy of a new benzodiazepine–dihydropyridine hybrid molecule (JM-20) in rodent models of ischemic stroke"

**The PDF file contains:**

Figure S1. Neuroscore of animals that completed the study (14-day survivors).

Figure S2. Neuroscore of animals that survived up to 7 days.

Figure S3. Neuroscore assigning the worst possible value (39) as an imputation method.

Figure S4. Neuroscore assigning the last available value as an imputation method.

Figure S5. Body weight gain expressed as a percentage of initial body weight (pre-surgery).

Figure S6. Corner test results 7 and 14 days after middle cerebral artery occlusion (MCAO), computed as the percentage of left turns relative to baseline data (pre-surgery day).

Table S1. Time to contact and time to remove in the Adhesive Removal Test and percentage use of the left paw in the Cylinder test (studies 1, 2, 4, 5).

Figure S7. Blood glucose levels before (A) and during (B) the post- middle cerebral artery occlusion (MCAO) period (Study 5).

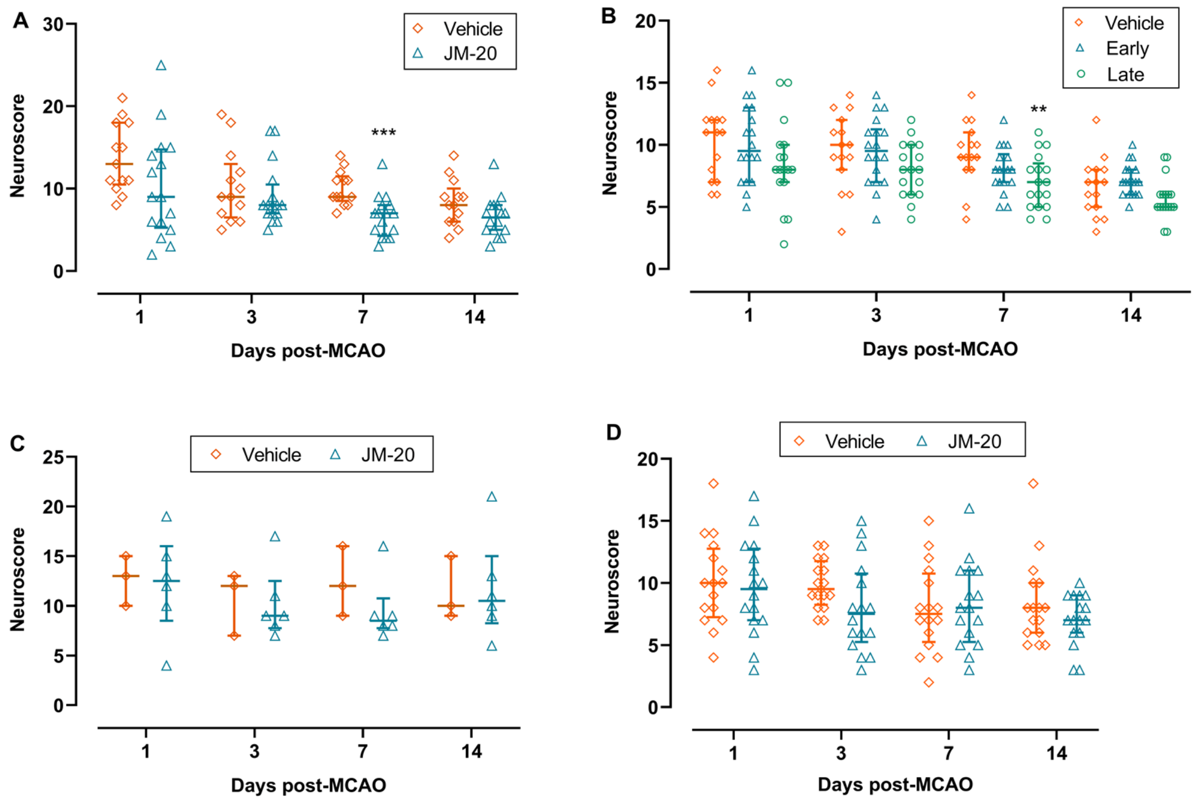


**Figure S1. Neuroscore of animals that completed the study (14-day survivors).** Median ± Interquartile range. (**A**) Data from Study 1 Havana 2019; ****P*<0.001 MCAO-Veh *vs*. MCAO-JM-20, by Unpaired *t*-test. (**B**) Data from Study 2 Berlin 2019; ***P*<0.015 MCAO-Veh *vs*. MCAO-JM-20 late treatment, by ordinary one-way ANOVA and Dunnett's multiple comparisons test. (**C**) Data from Study 4 Havana 2020; *P*>0.05, by Mann–Whitney U test. (**D**) Data from Study 5 Havana 2022; *P*>0.05, by Unpaired *t*-test.

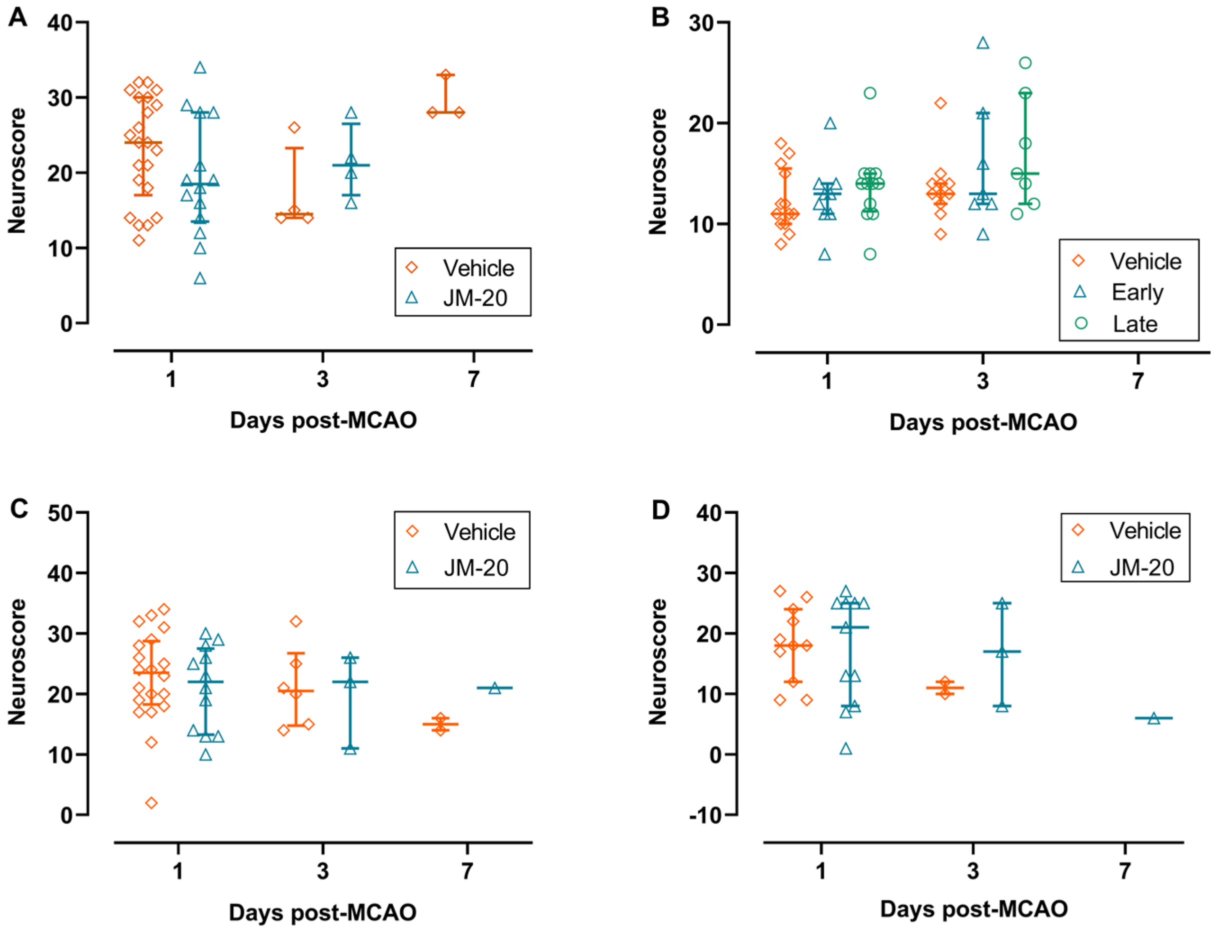


**Figure S2. Neuroscore of animals that survived up to 7 days.** Median ± Interquartile range. (**A**) Data from Study 1 Havana 2019. (**B**) Data from Study 2 Berlin 2019. (**C**) Data from Study 4 Havana 2020. (**D**) Data from Study 5 Havana 2022.

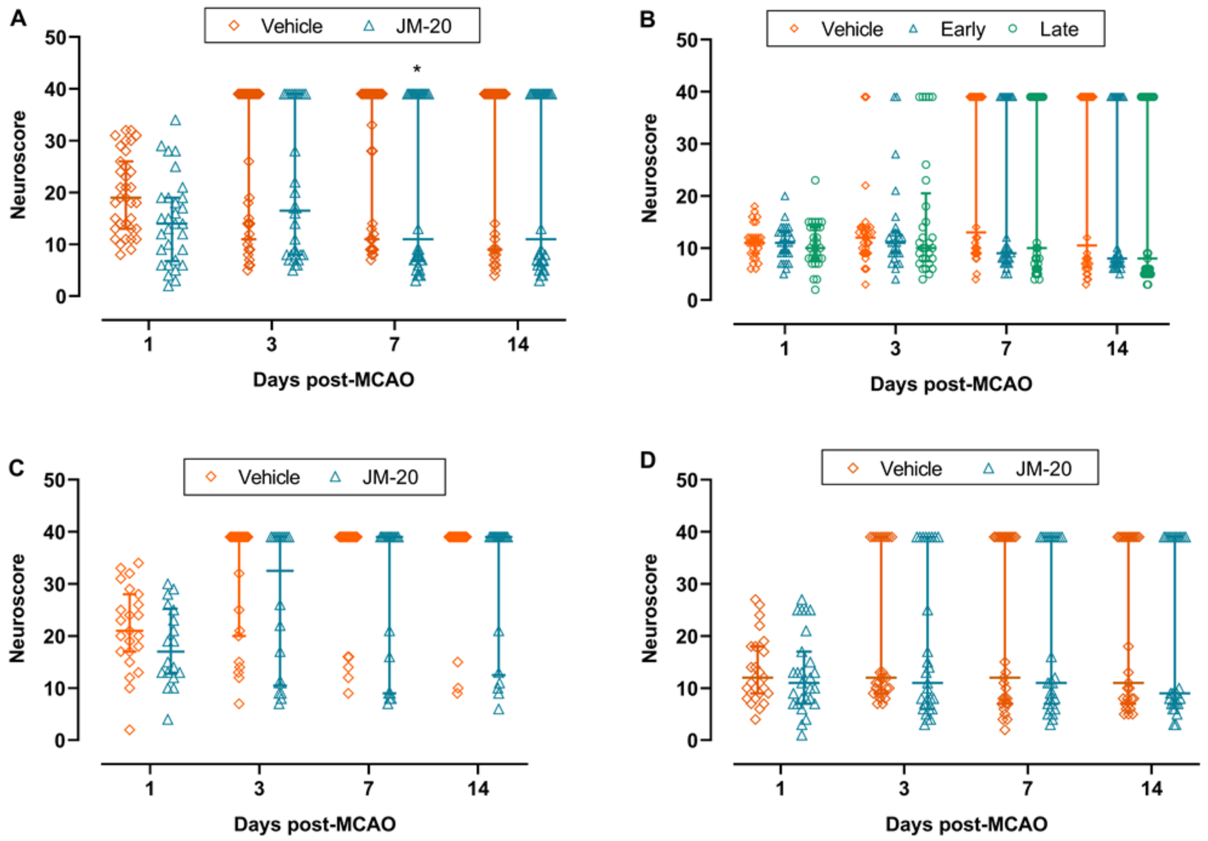


**Figure S3. Neuroscore assigning the worst possible value (39) as an imputation method.** Median ± Interquartile range. (**A**) Data from Study 1 Havana 2019. **P*<0.05 MCAO-Veh *vs*. MCAO-JM-20, by Mann–Whitney U test. (**B**) Data from Study 2 Berlin 2019. *P*>0.05, by Kruskal-Wallis test. (**C**) Data from Study 4 Havana 2020. *P*>0.05, by Mann–Whitney U test. (**D**) Data from Study 5 Havana 2022. *P*>0.05, by Mann–Whitney U test.

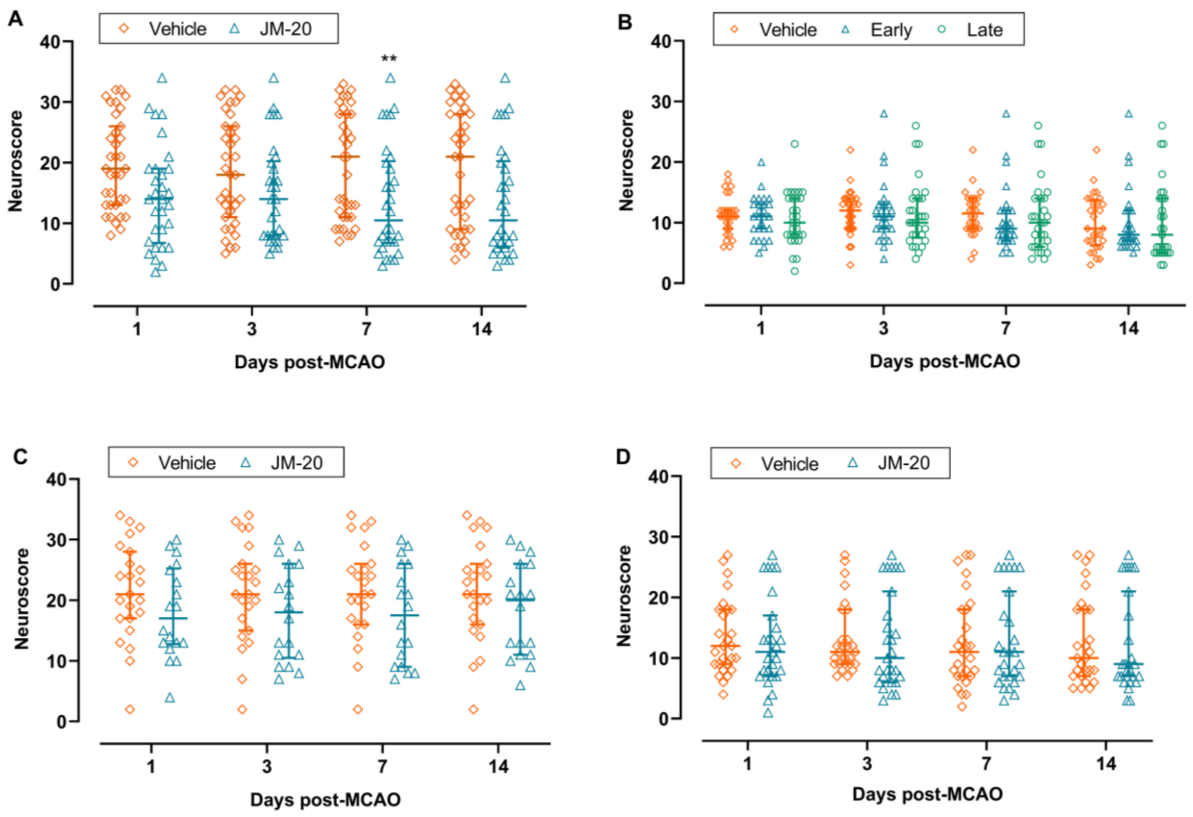


**Figure S4. Neuroscore assigning the last available value as an imputation method.** Median ± Interquartile range. (**A**) Data from Study 1 Havana 2019. ***P*<0.01 MCAO-Veh *vs*. MCAO-JM-20, by Mann–Whitney U Test. (**B**) Data from Study 2 Berlin 2019. *P*>0.05, by Kruskal-Wallis test. (**C**) Data from Study 4 Havana 2020. *P*>0.05, by Unpaired *t*-test. (**D**) Data from Study 5 Havana 2022. *P*>0.05, by Mann–Whitney U Test.

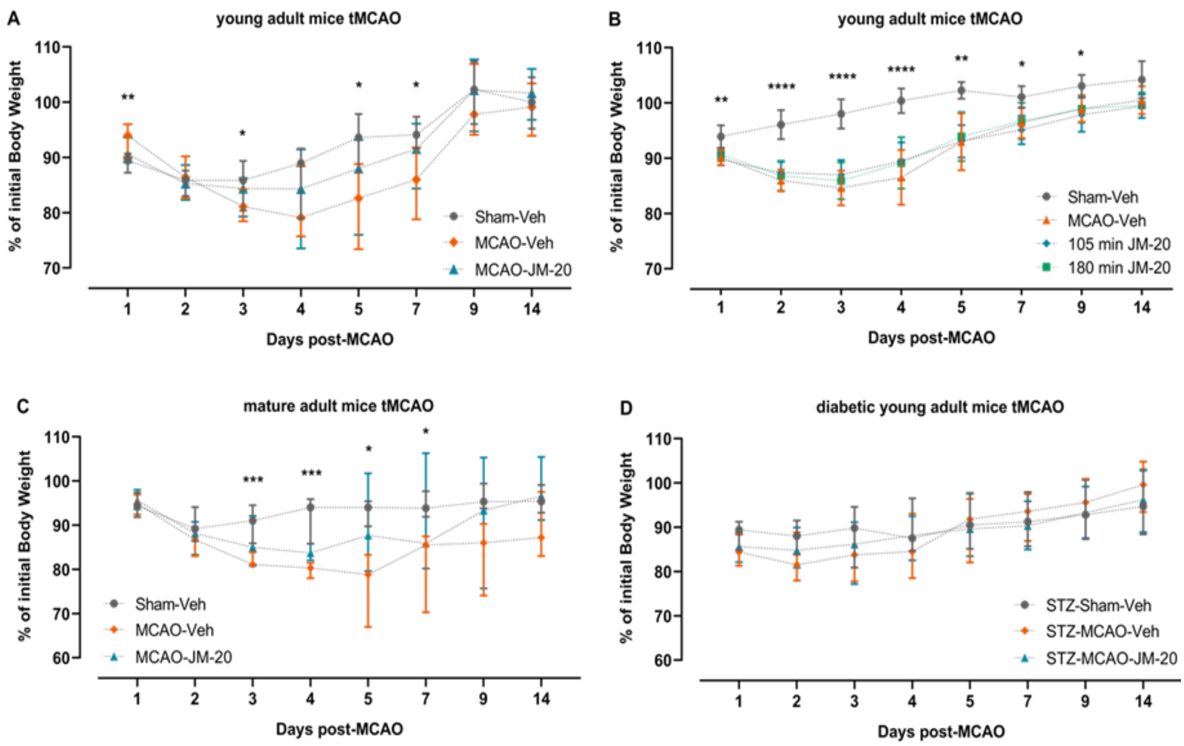


**Figure S5. Body weight gain expressed as a percentage of initial body weight (pre-surgery)**. Results are presented per individual study: (**A**) Study 1; (**B**) Study 2; (**C**) Study 4 and (**D**) Study 5. Data are Median ± Interquartile range. ****P<0.0001; ***P<0.001; **p<0.01; *p<0.05 *vs* MCAO-Veh group by Mixed-effects analysis and Dunnett's multiple comparisons test.

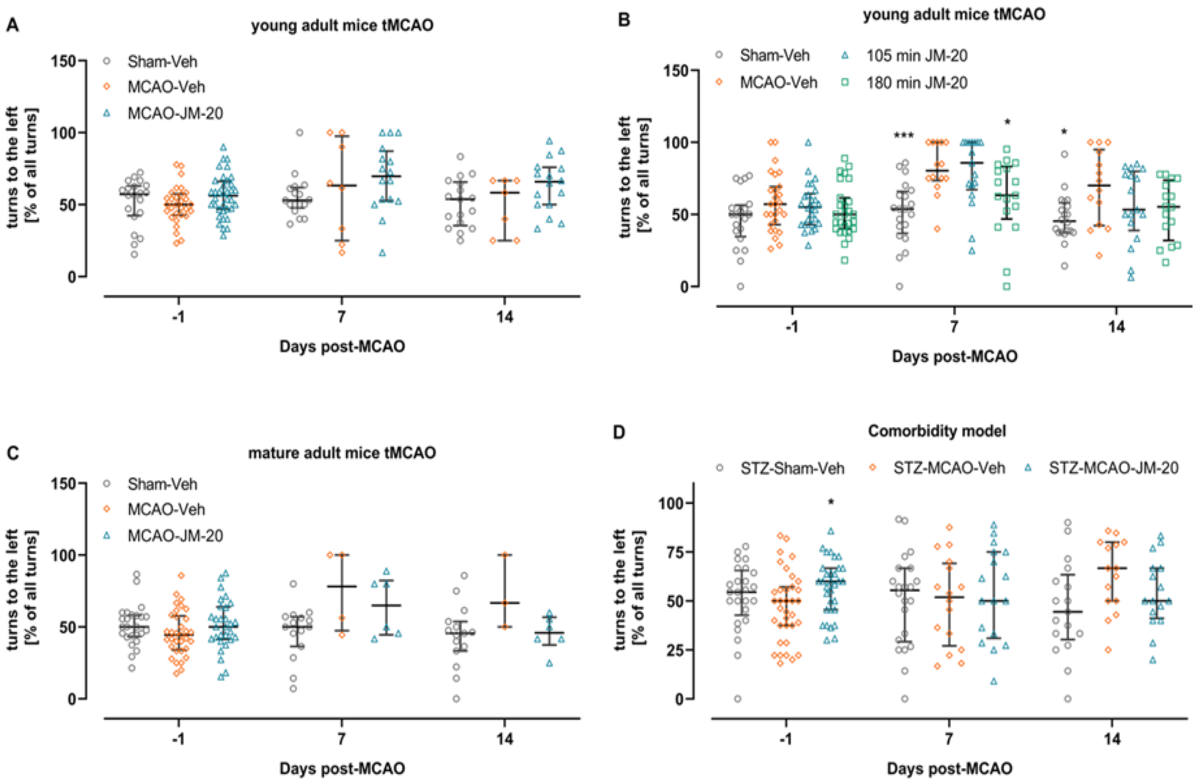


**Figure S6. Corner test results 7 and 14 days after middle cerebral artery occlusion (MCAO), computed as the percentage of left turns relative to baseline data (pre-surgery day).** (**A**) Study 1; (**B**) Study 2; (**C**) Study 4 and (**D**) Study 5. Data are Median ± Interquartile range. ***P<0.001; *P<0.05 *vs.* MCAO-Veh group by Mixed-effects analysis and Dunnett's multiple comparisons test.

**Table S1.** Time to contact and time to remove in the Adhesive Removal Test and percentage use of the left paw in the Cylinder test (studies 1, 2, 4, 5)

| Study #, location, year | Group | 1 week | | 2 week | | 2 week |
|---|---|---|---|---|---|---|
| | | Contact (s) | Remove (s) | Contact (s) | Remove (s) | Use of left paw (%) |
| 1. Havana 2019 | Sham-Veh | 41 ± 34.7 | 47 ± 35.9**** | 40 ± 37.8 | 46 ± 40.3 | 48 ± 6.1* |
| | MCAO-Veh | 46 ± 29.5 | 102 ± 29.7 | 36 ± 23.8 | 73 ± 41.3 | 59 ± 13.5 |
| | MCAO-JM-20 | 44 ± 30.1 | 101 ± 29.5 | 53 ± 34.2 | 99 ± 33.9 | 52 ± 11.7 |
| 2. Berlin 2019 | Sham-Veh | 10 ± 14.0 | 18 ± 16.5**** | 9 ± 12.2 | 21 ± 19.1*** | 35 ± 30.3 |
| | MCAO-Veh | 29 ± 28.7 | 85 ± 46.5 | 20 ± 28.0 | 69 ± 47.4 | 38 ± 31.8 |
| | MCAO-105 min JM-20 | 30 ± 27.9 | 106 ± 22.8 | 24 ± 24.1 | 84 ± 37.2 | 46 ± 25.3 |
| | MCAO-180 min JM-20 | 26 ± 34.3 | 85 ± 38.9 | 18 ± 16.8 | 74 ± 39.7 | 47 ± 30.3 |
| 4. Havana 2020 | Sham-Veh | 27 ± 18.9 | 32 ± 20.4 | 23 ± 26.7 | 27 ± 26.9 | 44 ± 15.5 |
| | MCAO-Veh | 37 ± 39.9 | 70 ± 47.9 | 9 ± 8.3 | 14 ± 8.8 | 52 ± 9.1 |
| | MCAO-JM-20 | 43 ± 22.6 | 67 ± 39.3 | 35 ± 29.0 | 55 ± 46.8 | 36 ± 19.6 |
| 5. Havana 2022 | Sham-Veh | 15 ± 13.3*** | 20 ± 13.6**** | 19 ± 16.5* | 26 ± 20.0*** | 48 ± 19.5 |
| | MCAO-Veh | 44 ± 34.3 | 70 ± 32.5 | 50 ± 38.3 | 82 ± 41.8 | 65 ± 15.3 |
| | MCAO-JM-20 | 37 ± 21.3 | 70 ± 34.8 | 40 ± 32.2 | 69 ± 41.5 | 65 ± 21.6 |

Data are shown as Mean ± Standard Deviation

*P<0.05; ***P<0.0001; ****P<0.0001 vs. MCAO-Vehicle group

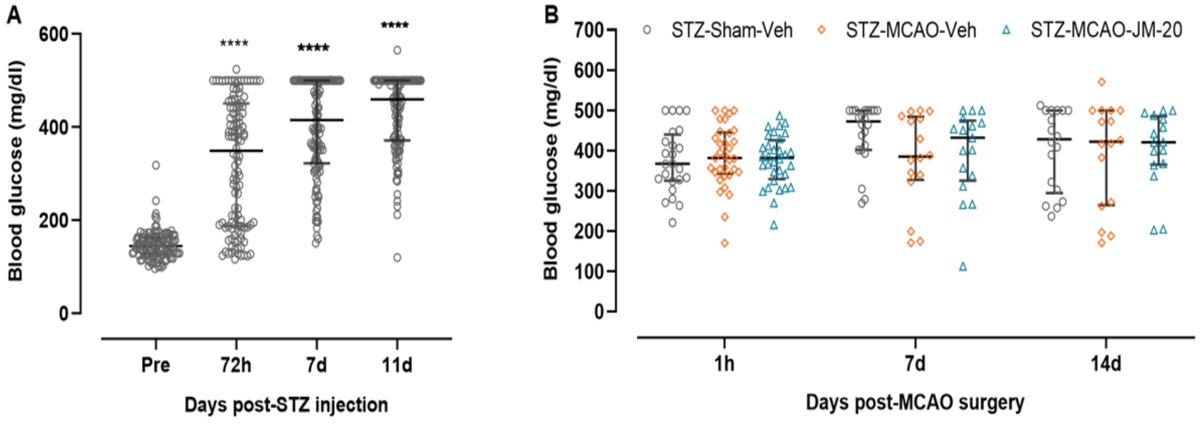


**Figure S7. Blood glucose levels before** (**A**) **and during** (**B**) **the post- middle cerebral artery occlusion (MCAO) period (Study 5)**. Streptozotocin (STZ)-induced hyperglycemia was confirmed in more than 80% of subjects and was maintained during the post-ischemic period (comorbidity model), regardless of the intervention (Sham/MCAO surgery) or treatment (vehicle/JM-20) received. Data are Median ± Interquartile range. ****P<0.0001 *vs.* pre-STZ injection, by Mixed-effects analysis and Dunnett's multiple comparisons test.
